# Precious3GPT: Multimodal Multi-Species Multi-Omics Multi-Tissue Transformer for Aging Research and Drug Discovery

**DOI:** 10.1101/2024.07.25.605062

**Authors:** Fedor Galkin, Vladimir Naumov, Stefan Pushkov, Denis Sidorenko, Anatoly Urban, Diana Zagirova, Khadija M. Alawi, Alex Aliper, Ruslan Gumerov, Aleksandr Kalashnikov, Sabina Mukba, Aleksandra Pogorelskaya, Feng Ren, Anastasia Shneyderman, Qiuqiong Tang, Deyong Xiao, Alexander Tyshkovskiy, Kejun Ying, Vadim N. Gladyshev, Alex Zhavoronkov

**Affiliations:** Insilico Medicine AI Limited, Abu Dhabi, UAE; Department of Medicine, Brigham and Women’s Hospital, Harvard Medical School, Boston, MA, USA

## Abstract

We present a multimodal multi-species multi-omics multi-tissue transformer for aging research and drug discovery capable of performing multiple tasks such as age prediction across species, target discovery, tissue, sex, and disease sample classification, drug sensitivity prediction, replication of omics response and prediction of biological and phenotypic response to compound treatment. This model combines textual, tabular, and knowledge graph-derived representations of biological experiments to provide insights into molecular-level biological processes. We demonstrate that P3GPT has developed an intuition for the interactions between compounds, pathologies, and gene regulation in the context of multiple species and tissues. In these areas, it outperforms existing LLMs and we highlight its utility in diverse case studies. P3GPT is a general model that may be used as a target identification tool, aging clock, digital laboratory, and scientific assistant. The model is intended as a community resource available open source as well as via a Discord server.

## Introduction

Biology has long been a discipline with a high affinity for machine-learning techniques. The discovery of DNA structure and advances in sequencing technologies have allowed scientists to treat molecular data as texts and thus apply various string-based and graph algorithms to study the evolution and regulation of cellular processes. We have witnessed the emergence of an interdisciplinary research field founded on the assumptions that living systems can be described as mathematical abstractions and that their behavior can be accurately predicted with increasingly powerful compute.

Another milestone occurred in the 1990s, when the scientific community achieved the feat of sequencing whole genomes. With the advent of shotgun-sequencing techniques, the rate of biodata generation increased dramatically, and online public repositories, such as the Gene Expression Omnibus (GEO), were founded; these repositories remain the field’s most essential web resources^1^. Open access to large amounts of data perfectly positioned scientists to contribute to the upcoming revolution in data science and resulted in the quick adoption of breakthrough approaches developed in other fields.

The progress in machine-learning approaches culminated in the neural network boom of the 2010s. The variety of available architectures, combined with the inherent ability to capture nonlinear dependencies, secured neural networks’ position as the algorithms of choice for solving any tasks involving big data. In biomedical research, neural networks have been used to analyze imaging data, develop novel drugs, apply biological knowledge graphs (KGs), study aging processes, annotate genomes, predict protein structures, and even emulate living organisms.

Compared to other machine-learning methods, neural networks demonstrate outstanding performance in these tasks, occasionally outperforming trained humans. However, these models have been trained to carry out highly specific tasks and thus can hardly be called artificial intelligence (AI). The high level of specialization achieved in these models also poses a significant challenge to their wider adoption, since their range of application may be limited to the various technicalities associated with sample collection, sample preparation, and—in many cases—data processing protocols, which are likely to introduce bias into the data produced. Effectively, data obtained through different means may represent entirely separate domains, resulting in a reproducibility crisis^2^.

Omics studies are particularly susceptible to this issue. Despite extensive research on normalization and harmonization methods, cross-platform data aggregation remains an unresolved challenge. As new platforms emerge and older ones are cycled out of production, the problem is further aggravated, as specialized AI models trained with a particular format in mind are rendered useless once that format is abandoned.

One solution to the reproducibility crisis in biomedical research is organizational. It involves the formation of consortia to manage end-to-end data acquisition and preserve biospecimens for follow-up measurements. Such long-lasting projects include NHANES, the UK Biobank, LINCS, the Framingham Heart Study, and other biobanks with a long history of meticulous data management procedures, which ensures research continuity.

Another approach involves the development of more general AI systems that are resistant to technical biases and can operate at a sufficiently high level of abstraction to extract and combine knowledge from various sources. The transformer architecture developed by Google in 2017 rapidly became the industry standard for solving tasks characterized by high uncertainty and a lack of formal definition^3^. Successfully leveraging the assumption that any problem and its answer can be expressed in natural language, generative pretrained transformers (GPTs) took the world by storm in 2022 with the release of ChatGPT. Most recently, base models fine-tuned to specific research areas have seen considerable development. In the same year, BioGPT, which was pretrained on a corpus of 15 million PubMed texts, was released. It was the first such technology able to effectively carry out relation extraction, classify scientific documents, and answer biomedical questions^4^. Another notable instance is OpenBioLLM, released in 2024, which outperforms OpenAI’s GPT-4 in a number of benchmarks and can be used to parse clinically relevant information and answer biomedical questions^5^.

While these models have demonstrated stellar performance in conveying biomedical information and providing evidence-based answers, their functionality is greatly constrained by the limitations of the type of textual data used. The information made available to these GPT-based models is limited to the highly distilled research output seen in academic publications. Meanwhile, the original data on which these research projects are based remain hidden from the model, regardless of any unextracted knowledge remaining in the data. Arguably, specialized large language models (LLMs) suffer from a self-imposed handicap and lack the ability to provide novel hypotheses not yet stated or implied in the literature^6^.

In this article, we present Precious3-GPT (P3GPT), a new type of biomedical LLM trained on multimodal data consisting of PubMed publications, KGs, and—most importantly—tabular data representing unadulterated research information yet to be interpreted (**Figure 1**). By enabling it with multimodal capabilities, we have ensured that it can learn from a much wider range of resources than other LLMs. P3GPT has learned to reproduce the raw output of research experiments and can thus operate at a lower level than other biomedical LLMs. Its native output format is expressed in terms of genes and chemical compounds, effectively making P3GPT a digital lab suitable for various *in vivo*, *in vitro*, or clinical experiments. We showcase P3GPT’s abilities in a range of aging-related, biomedical entity classification, and omics domain tasks.

**Figure 1.**
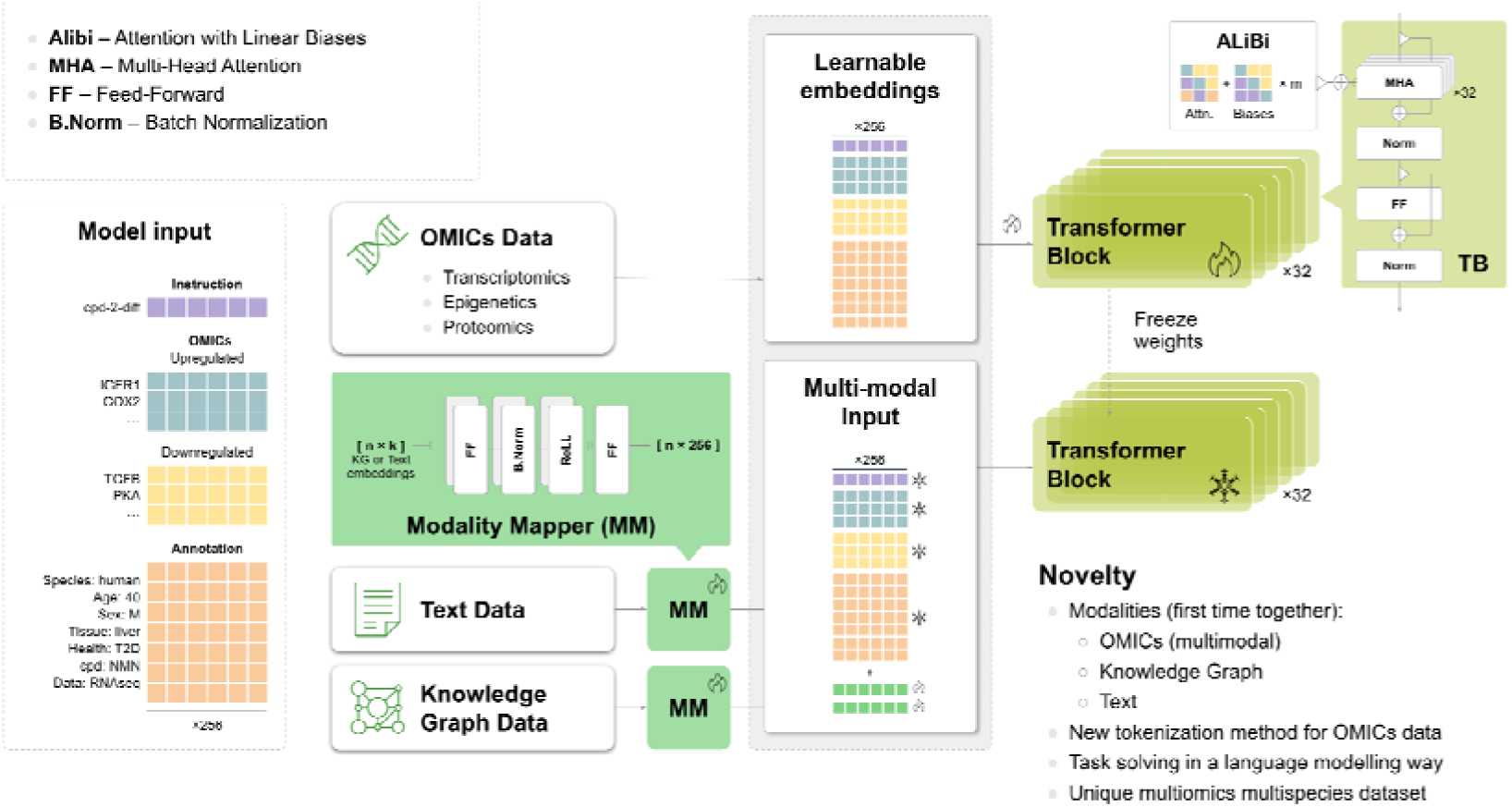
P3GPT features a novel architecture enabling efficient omics-data training and multi-modal extensions. Omics tabular observations are transformed into structured input prompts to train the transformer block. Additional modalities (text and knowledge graph) are embedded through modality mapper units as extensions to the frozen transformer block. See more in Methods.

The range of research settings it supports and its efficient learning rate make P3GPT a unique model that holds the potential to become a centerpiece of a fully-automated, AI-managed research laboratory. With the help of autonomous AI agent frameworks, LLMs can carry out functions beyond text generation and parsing to plan experiments, interpret data, modify existing scripts to support new packages and data formats, and basically support any research task provided the tools (**Figure 2**). As an LLM, P3GPT is ready to be implemented as such a tool, which we demonstrate by using its output in a real-life *in vitro* experiment aimed at identifying novel geroprotectors.

**Figure 2.**
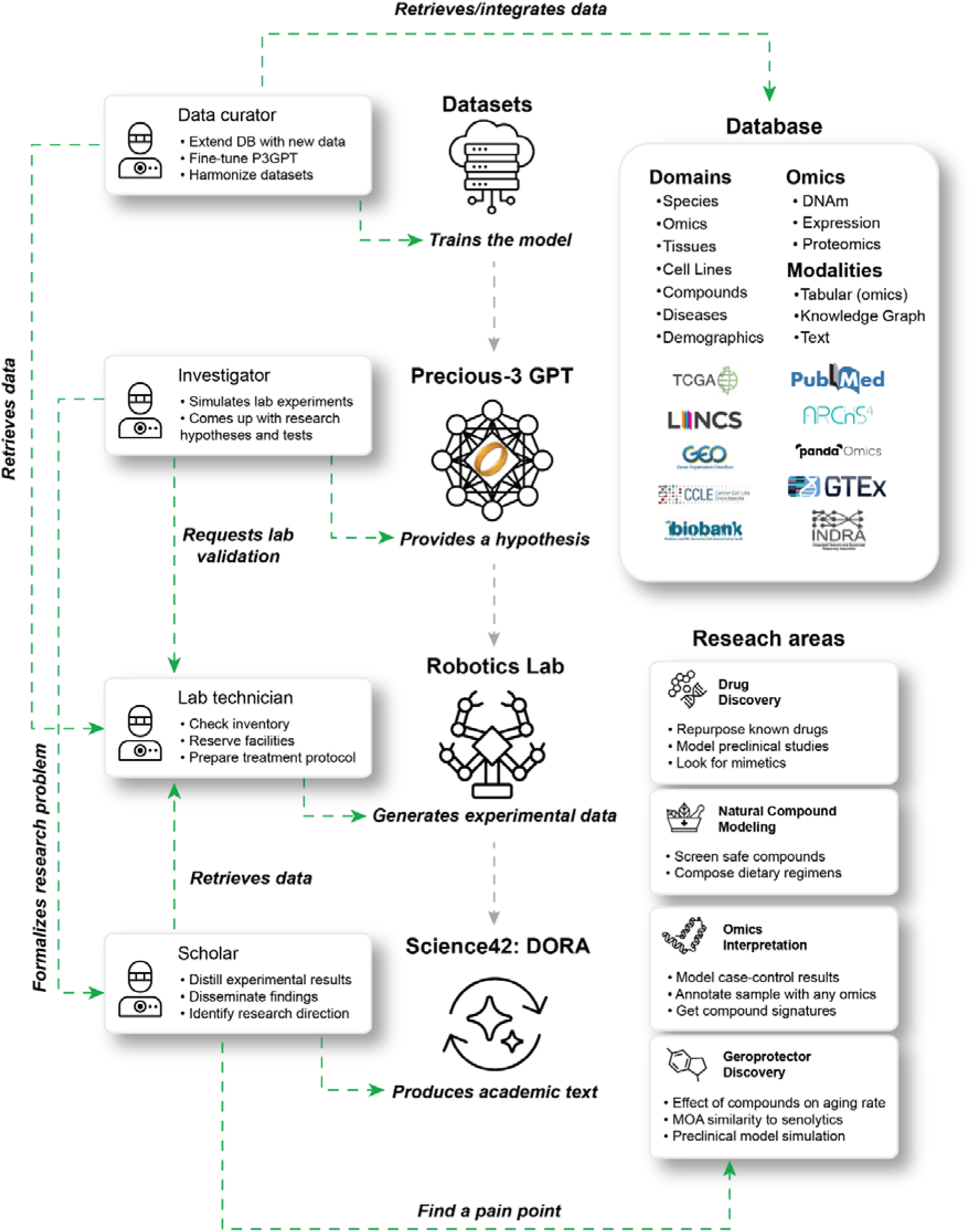
P3GPT’s ability to operate with omics data enables it to serve as a hypothesis engine in an autonomous multi-agent system tasked with conducting biomedical research. Autonomous agents (left column) may use web services (center) as tools to carry out the functions typical for a biomedical research group. In such a collective of AI agents, P3GPT serves as a fast and affordable way to screen compounds or run omics experiments to generate hypotheses, e.g. “maslinic acid is a promising senolytic”. Other agents act on this hypothesis preparing a lab protocol and scheduling in vitro validation. Its results are shared with agents who can digest the findings in natural language for a human-assisted review. After the review, the results are integrated into the ever-growing database to be used for P3GPT fine-tuning. Finally, P3GPT is ready to generate new hypotheses, thus closing the loop.

## Results

### Data collection and training

To ensure that P3GPT developed an understanding of the interconnectedness of biological entities in the context of omics, we aggregated high-level, uninterpreted data from various omics experiments involving gene expression, DNA methylation, and proteomics. After applying extensive quality control procedures (see Methods), our data amounted to over 1.2 million observations featuring 63,376 biological entities (**Tables 1–2**). This data collection represents a wide range of experimental settings, such as cross-sectional human studies, murine case-control, and *in vitro* chemical intervention studies.

**Table 1.** P3GPT can interpret >60,000 biochemical entities, such as genes, compounds, and tissues. Overview of biological entities that P3GPT can interpret in the context of omics experiments. The model treats each entity as a distinct token that provides the context for an experiment or is generated through inference. The model also features 74 utility tokens that serve as tags and instructions.

| Biological Entity | N entities | Entity Examples |
| --- | --- | --- |
| Gene | 25,332 | ISG15, CXCL2 |
| Compound | 22,241 | BRD-K32665240, Floxuridin |
| Pathway | 13,439 | GOBP: Recognition of apoptotic cell |
| Mechanism of action | 661 | Renin Inhibitor |
| Condition | 635 | MONDO:0017396, EFO:0000305 |
| Age & Age groups | 394 | 21, 30-40 |
| Tissue | 300 | Heart, Joint |
| Cell Line | 269 | HT-29, IMR90 |
| Dose | 76 | 0.12uM, 200uM |
| Time of exposure | 15 | 96h, 8d |
| Data Type | 9 | Proteomics, RNAseq |
| Species | 3 | Homo sapiens |
| Gender | 2 | Male, Female |

**Table 2.**
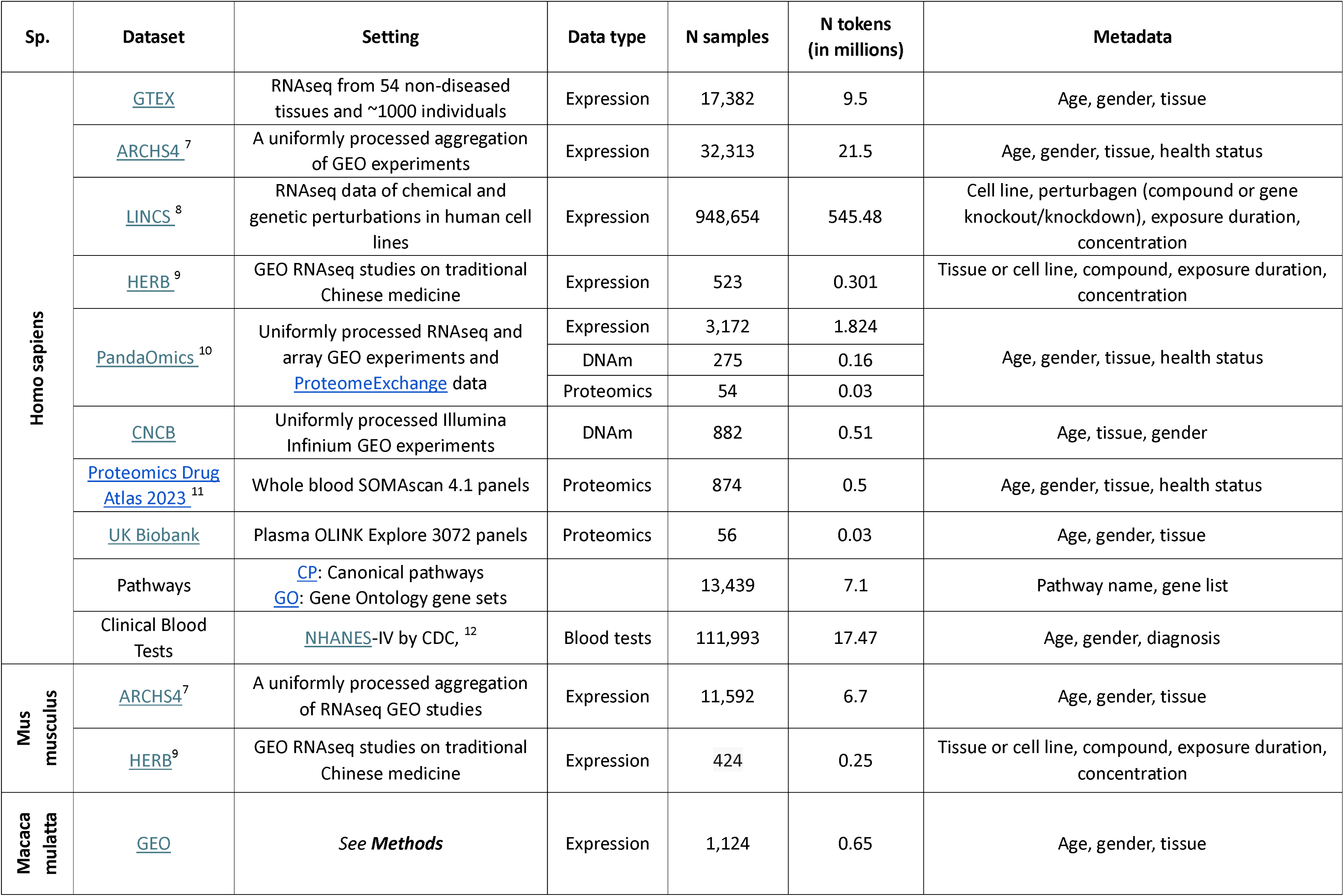
P3GPT was trained on a collection of 1.2 mln omics samples. The data was collected from publicly available resources featuring experiments with transcriptomic, epigenetic, or proteomic pr ofiles, such as GEO, LINCS and others. We also expanded the training set with gene lists organized in ontologies, such as Gene Ontology (GO), and clinical blood tests from NHANES to enable more use cases in the future

In our dataset, 18,224 human samples are annotated with age metadata and represent all age groups, including embryos and supercentenarians (**Figure 3A**). Similarly, our dataset contains 23,024 murine and 1,124 simian samples with known chronological age. The most represented tissues in our aggregate dataset include breast, prostate, lung, kidney, and skin, which together account for >380,000 observations (**Figure 3B**).

**Figure 3.**
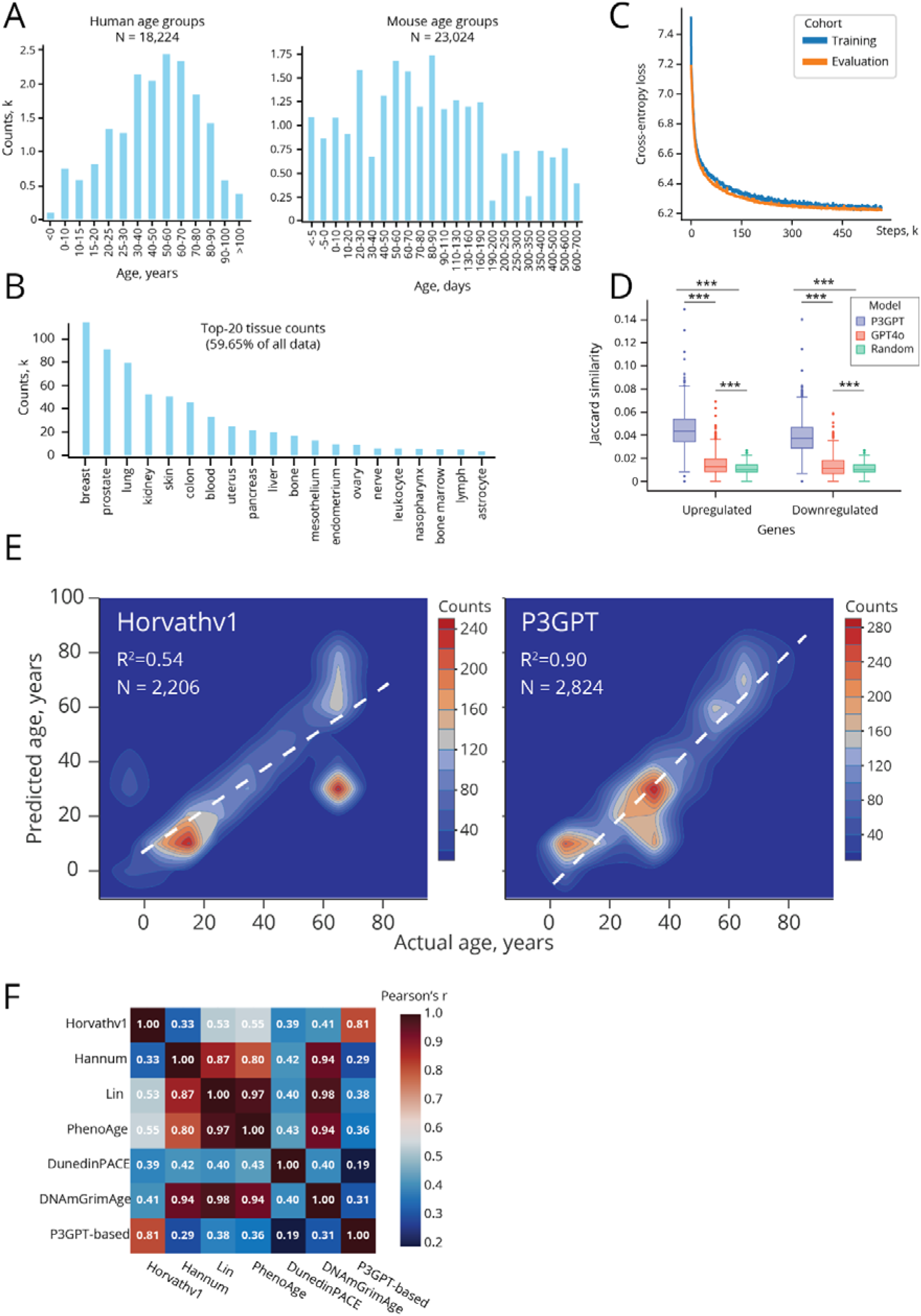
P3GPT was trained on data from multiple species, tissues, and age groups to simulate settings such as aging studies and chemical screenings. (A) P3GPT was trained on data from all age groups in both humans (left) and mice (right), spanning embryos and old individuals. (B) P3GPT’s training set features a wide selection of tissues to grant it tissue-context capability. (C) P3GPT’s TB learning curve demonstrates an efficient convergence of the training process. The training was arrested to add KG and text modalities at 566k steps, when the loss stopped decreasing. One step constitutes 12 samples, processed with 4 GPUs. (D) P3GPT outperforms ChatGPT-4o in the task of predicting the effect of a compound on a cell line in a subset of 805 LINCS entries not used in training. The Jaccard similarity of the P3GPT-generated perturbation signatures is 3–4 times higher than that of the GPT4o-generated signatures. *** — Mann–Whitney U-test P-value < 0.001 (E) P3GPT can be used to train an aging clock using the averaged embeddings of the most methylated gene promoters. The P3GPT-based predictor outperformed Horvath’s aging clock on a validation set of 2,824 human blood DNAm samples. (F) The P3GPT-derived aging clock offers a unique perspective on aging. Predictions of a P3GPT-based aging clock show high correlation with those of a multi-tissue Horvath’s 2013 clock but weaker correlations with blood-based DNAm aging clocks. Horvath’s 2013 clock R^2^=0.54; P3GPT-based clock R^2^=0.90.

Omics tabular data were transformed into structured input prompts containing the lists of the most and least abundant omics features, metavariable. The prompts were then supplemented with instructions to denote the experimental context and submitted to training the transformer block (TB). The training continued for 566,000 steps with 48 samples per step, at which point the cross-entropy loss function plateaued (**Figure 3C**).

In addition to tabular omics data, the training set included text and KG data, represented by GenePT-derived embeddings and heterogeneous graph transformer embeddings of 25,332 genes. After the omics training stage, TB’s weights were frozen and modality mapper (MM) units were trained to extend the learned prompt embeddings. The MM training continued for 2,350 steps with 24 samples per step until the loss function reached a plateau.

The full list of settings used in training P3GPT is described in Methods, and P3GPT’s architecture is shown in **Figure 1**.

### Gene list generation

One of the research settings P3GPT is trained to replicate is a chemical screening with omics measurements. To illustrate P3GPT’s competence in this core task, we employed P3GPT to generate lists of differentially expressed genes (DEGs) for a holdout set of compound screening experiments.

The holdout set featured 805 observations of chemically induced expression perturbations across 582 compounds and 102 cell lines. P3GPT was instructed to return DEG lists of the same length as in the original data: 100 or 250 items long. The Jaccard similarity between the P3GPT-produced and the original DEGs was 0.0387 for downregulated genes and 0.0447 for upregulated genes—values that are 3.49–3.98 times higher than those achieved by random lists and 2.92–3.06 times higher than those of GPT4o-generated DEGs (Jaccard similarity = 0.0132–0.0146, **Figure 3D**).

Despite a substantial difference in the number of parameters and the scale of the training set, P3GPT outperforms GPT4o in the task of predicting the response to a chemical perturbation. We see this as proof of P3GPT’s multimodal architecture superiority in the context of omics-level research, compared to text-trained models.

### P3GPT entities are associated with biological processes

If P3GPT may produce biologically relevant data, we may also expect its embeddings to represent experimentally validated annotations in terms of protein localization and metabolic pathway engagement. To test this hypothesis, we extracted P3GPT’s embeddings for all 25,332 human genes it could interpret to train binary classifiers for 18 high-level Gene Ontology (GO) terms randomly selected among those containing >100 genes.

The derived classifiers proved to be efficient (ROC-AUC [1 [0.743–0.877]) in annotating all three aspects of genes as defined in GO: molecular functions, biological processes, and cellular components.

Using the agent-based validation pipeline, we then replicated the same workflow with four other LLMs designed specifically to solve biomedical problems, including BioGPT and OpenBioLLM (**Table 3**). By comparing the results for this array of LLMs, we concluded that the P3GPT-derived representations are the most descriptive of the genes’ involvement in biological processes and protein localization. Thus, it is expected to be a more reliable tool than text-trained biomedical LLMs for the purposes of target identification and drug discovery, both settings that require careful consideration of how proteins and compounds interact with each other to affect regulatory pathways.

**Table 3.**
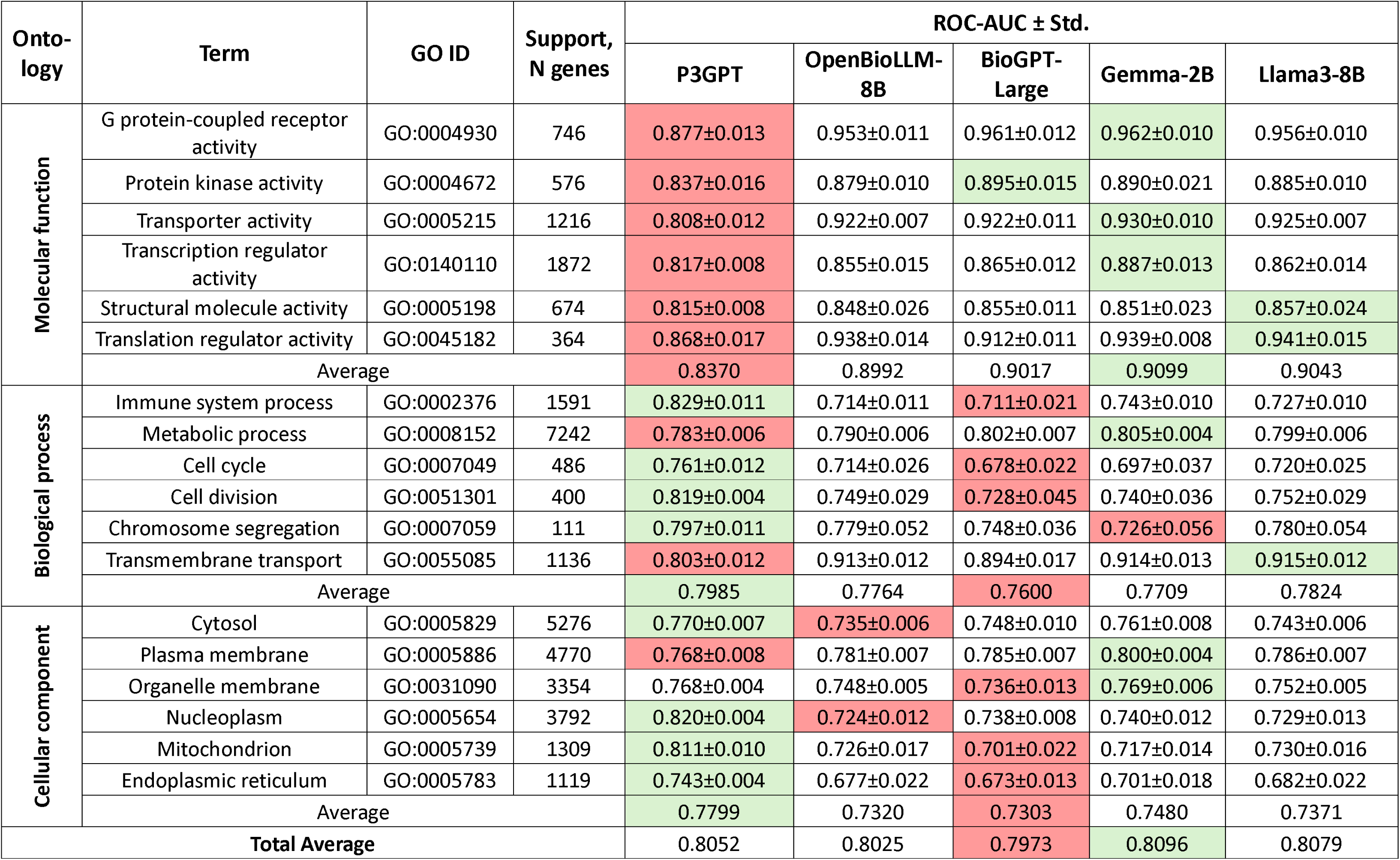
P3GPT outperforms much larger LLMs in the task of annotating genes with their biological function and cell localization. Performance of Gene Ontology (GO) term classifiers trained on embeddings extracted from various LLMs. The P3GPT-based classifiers exhibited superior performance with 8 out of 18 high-level GO terms but underperformed in classifying the genes’ molecular functions. ROC-AUC — area under the receiver operating curve; Std — standard deviation. The background set consisted of 22,509 human genes. The green and red highlights denote the best- and worst-performing classifiers for each term

### P3GPT-derived aging clocks

To illustrate the utility of P3GPT in aging research, we tested it as an age predictor trained on a collection of 10,750 blood samples from public DNAm profiling GEO experiments. To build the clock, we first averaged P3GPT embeddings of the 50 most methylated genes for each sample to obtain 360 dimensions-long vectors. Then we stacked the embeddings with a matrix of average ß-values on the promoters of 360 genes selected as the features with the highest SHAP importance scores in a CatBoost model trained on the ß-values of all 25,332 genes.

The stacked P3GPT embeddings and ß-values were then used to train a CatBoost predictor of chronological age, which we tested in a validation set of 2,824 samples. The resulting aging clock displayed a mean absolute error of 4.78 years and R^2^=0.90 (**Figure 3E**). Additionally, we applied SHAP value feature importance analysis to assess the degree of P3GPT’s embeddings to the accuracy of the predictor. The cumulative importance of the P3GPT features accounted for 12.2% of total importance. We thus show that P3GPT-derived features have proven to be a measurable source of additional aging-related information that can improve the performance of an aging clock. In other words, P3GPT has successfully internalized the connection between gene methylation levels and aging, which can be exploited in applications building on top of P3GPT.

To frame our P3GPT-based clock in the context of other aging clocks, we utilized the predictions for our validation set from the ClockBase platform ^13,14^. We then inspected the correlations between all pairs of six commonly used aging clocks and our model’s predictions. The P3GPT-based clock was highly concordant only with the Horvath’s 2013 clock (Pearson’s r = 0.81), which is a multi-tissue biomarker. It showed lower correlations coefficients with blood-based clocks (Hannum ^15^, GrimAge ^16^, PhenoAge ^17^, Lin ^18^) as well as with the biomarker (DunedinPACE ^19^) that assesses the rate of aging, which is also based on blood (Figure 3F). Thus, P3GPT can derive reliable multi-tissue biomarkers of chronological age.

We also instructed the in-house multi-agent system to apply a similar methodology to train a pan-mammalian DNAm aging clock using the GEO dataset featuring DNAm samples from 348 species ^20^. To enable compatibility with P3GPT, the CpG sites in unseen mammalian species were mapped onto human homologs, and a regressor was trained to predict the chronological age in each animal. The accuracy of the predictor ensemble was marked by a 10.06% mean absolute percentage error (MAPE), an R^2^ value of 0.68, and a Pearson’s r value of 0.83 in a test set of 1,406 multi-species multi-tissue samples (**Figure 4A**). Thus, P3GPT showed its potential utility in studying aging not only in humans, but across the mammalian phylum in general.

**Figure 4.**
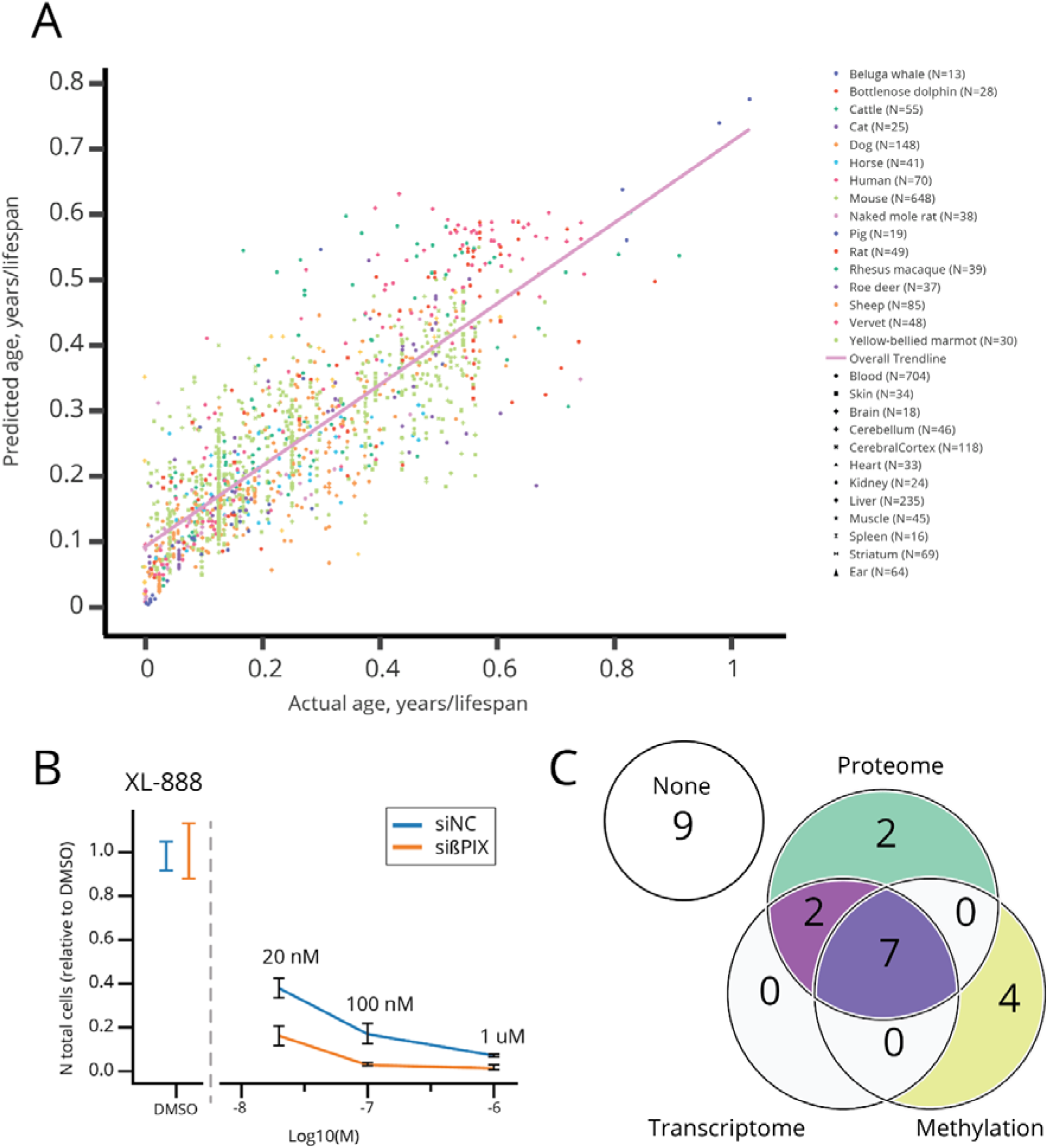
P3GPT shows unique capabilities, such as age prediction in unseen species, discovery of novel geroprotectors, and cross-omics target discovery. (A) P3GPT’s gene embeddings can be used to train a pan-mammalian aging clock, even in species not encountered in training. A chronological age regressor was trained for each species using stacked embeddings of the top 500 hypermethylated human homologs. The ensemble shows R ^2^ = 0.68 across all subsets. (B) XL-888 is a potential senolytic, as identified by P3GPT. It shows selective cytotoxicity to senescent cells in an IMR-90 sißPIX induced senescence model with a more pronounced senolytic effect in low nanomolar concentrations. (C) P3GPT can reliably detect clinical targets with any of its omics modalities. Clinical targets for 15 out of 24 diseases were present in P3GPT generations with corresponding experimental conditions. Furthermore, the target genes for 7 indications appeared in all three available generation modalities: proteomic, transcriptomic, and methylomic.

### P3GPT-derived geroprotectors

To assess P3GPT’s practical utility, we used it as a hypothesis generator in an *in vitro* anti-aging experiment. To obtain a list of potential geroprotectors from P3GPT, we sequentially executed it with two instructions. First, we applied the <age2diff> instruction to obtain expression signatures differentiating younger (20 years) and older (80 years) adults. Second, we applied the <diff2cpd> instruction to generate the compounds expected to reverse the signature identified in the first step in IMR90 cells. Upon manual curation to exclude toxic and commercially unavailable compounds, the 22 molecules listed in **Table 4** were selected for screening in an *in vitro* senescence model (see **Methods**).

**Table 4.** P3GPT identified 22 compounds as potential geroprotectors that were examined in IMR90 cells. A total of 54 dilutions of 22 unique molecules were assessed in a ßPIX-siRNA senescence model with ßGal staining as a senescence marker. CVD — cardiovascular disease, HRT — hormone replacement therapy.

| Compound | CHEMBL ID | Concentrations tested | Known indications |
| --- | --- | --- | --- |
| BI-2536 | CHEMBL513909 | 10 uM | Neoplasms (phase-2) |
| Bisindolylmaleimide-IX | CHEMBL6291 | 10 uM | Leukemia (phase-2) <sup>21</sup> |
| Bortezomib | CHEMBL325041 | 20nM, 100nM, 1uM, 10uM, 20uM | Mantle cell lymphoma, multiple myeloma |
| Celastrol | CHEMBL301982 | 10uM | Neoplasms (preclinical), CVD (preclinical) <sup>22–24</sup> |
| Clomethiazole | CHEMBL315795 | 20nM, 100nM, 1uM, 10 uM | Sedative |
| CYT-997 (Lexibulin) | CHEMBL552212 | 10uM | Multiple myeloma (phase-2) |
| Dapsone | CHEMBL1043 | 20nM, 100nM, 1uM, 10uM, 20uM | Acne, leprosy |
| Estradiol cypionate | CHEMBL1200973 | 20nM, 100nM, 1uM, 10 uM | HRT in menopause |
| GMX-1778 | CHEMBL17289 | 10 uM, 20uM | Neoplasms (phase-1) |
| Indisulam | CHEMBL77517 | 1 uM | Neoplasms (phase-2) |
| Ixazomib | CHEMBL2141296 | 10 uM | Neoplasms |
| Lucitanib | CHEMBL2220486 | 20nM, 100nM, 1uM, 10 uM | Neoplasms (phase-2) |
| Maslinic acid | CHEMBL201515 | 10 uM, 20uM | NA |
| Metronidazole | CHEMBL137 | 20nM, 100nM, 1uM, 10 uM | Infections, pneumonia |
| MRS-1754 | CHEMBL273807 | 20nM, 100nM, 1uM, 10 uM | CVD <sup>25</sup> |
| NVP-AUY922 (Luminespib) | CHEMBL252164 | 10uM | Neoplasms (phase-2) |
| Obatoclax | CHEMBL408194 | 10uM | Neoplasms (phase-3) |
| Pidotimod | CHEMBL1488165 | 20nM, 100nM, 1uM, 10 uM | Asthma and other respiratory diseases <sup>26,27</sup> |
| Sonidegib | CHEMBL2105737 | 10uM, 20uM | Basal cell carcinoma |
| TP-0903 | CHEMBL2022968 | 10uM | Neoplasms (phase-2) |
| Triptolide | CHEMBL463763 | 10uM | Polycystic kidney disease (phase-3) |
| XL-888 | CHEMBL4297451 | 20nM, 100nM, 1uM, 10uM | Skin cancer (phase-1) |
| Rapamycin | CHEMBL413 | 50nM | Known senomorphic |
| Navitoclax (ABT-263) | CHEMBL443684 | 1uM | Known senolytic |

The selected compounds were tested in 1-4 concentrations (10 nM-20 uM) to assess their ability to reduce the number of ß-galactosidase (ßGal)-positive IMR90 cells subjected to sißPIX-induced senescence. To affirm the validity of the used model, we used two control compounds. Rapamycin was selected as a positive control of senomorphic activity — a lower senescent cell portion in compound-treated sißPIX cultures, compared to untreated sißPIX cultures. Navitoclax (ABT-263) was selected as a positive control of senolytic activity — a lower cell survival rate in compound-treated sißPIX cultures, compared to compound-treated non-senescent cultures.

Among the 22 screened compounds, 5 showed a significant (p-value < 0.05) senomorphic potential, resulting in a hit rate of 23% (**Table 5**). Compared to the reference senomorphic (50 nM rapamycin), the candidate geroprotectors identified are less potent, with only 3 compounds reducing the number of ßGal cells by > 5%: maslinic acid (20 uM), estradiol cypionate (10 uM), and dapsone (20 uM). Two more compounds can be added by applying a more relaxed p-value threshold (= 0.10): MRS-1754 and clomethiazole. The list of potential geroprotectors can be further extended to 8 entries if cytotoxic compounds, such as XL-888, are considered (**Figure 4B**).

**Table 5.** P3GPT achieved a 23% hit rate in the task of identifying non-toxic geroprotectors for a custom setting. In total, eight compounds proposed by P3GPT and included in the screening exhibited potential senolytic or senomorphic activity activity. For five compounds out 22 considered, no cytotoxicity towards non-senescent cells was detected. Senolytic action is defined as the change in cell survival between compound-treated senescent and non-senescent cultures. Senomorphic action is defined as the change in ßGal+ cells between compound-treated and untreated senescent cultures. Cytotoxicity is defined as the change in cell survival between compound-treated and untreated non-senescent cultures. (see Methods) NS ̶ not significant (P-value > 0.10), * ̶ P-value ≤ 0.10; ** ̶ P-value < 0.05. All dilutions were observed in three replicates. Rapamycin and navitoclax were used as the reference senomorphic and senolytic compounds, respectively.

| **Compound** | **Selected concentration** | **Senolytic action, %** | **Senomorphic action, %** | **Cytotoxicity, %** |
| --- | --- | --- | --- | --- |
| Clomethiazole | 100 nM | NS | -2.53 ** | NS |
| Dapsone | 20 uM | >0* | -5.01 ** | NS |
| Estradiol cypionate | 10 uM | NS | -5.20 ** | NS |
| Maslinic acid | 20 uM | NS | -5.66 ** | >0** |
| Metronidazole | 1 uM | -9.4 * | -3.14 ** | >0** |
| MRS-1754 | 100 nM | >0* | -5.41 ** | >0** |
| Pidotimod | 10 uM | NS | -3.09 ** | >0** |
| XL-888 | 20 nM | -21.26 * | >0** | -62.3 * |
| Rapamycin | 50 nM | >0* | -10.31 ** | NS |
| Navitoclax (ABT-263) | 1 uM | -98.4 * | -12.10 ** | NS |

This experiment confirms P3GPT’s ability to serve as a biomedical hypothesis generator, realistically simulate the aging process, and propose novel potential geroprotectors, whose effects are validated *in vitro* with a high success rate.

### Multimodal target enrichment

P3GPT was trained on biodata modalities representing different levels of regulation, including gene expression, DNA methylation, protein levels and protein interactions. To validate that the model successfully internalized entity relations across all omics levels, we implemented the following target identification test. First, we prepared a collection of 24 diseases that affect different tissues and instructed P3GPT to generate differentially methylated, expressed, and translated genes. The conditions in this experiment were selected based on a large number of known target genes (>10) and a sufficiently large representation of the affected tissue in the training set (see Methods).

We then counted the number of actual clinical target genes in the 300 likeliest output tokens in each indication–modality combination. We identified 15 indications whose output tokens were significantly (P-value < 0.05) enriched for real targets for differential expression, among which 7 were enriched for clinical targets in all three omics modalities (**Figure 4C**).

This experiment demonstrates that the proteomic, epigenetic, and transcriptomic modes of P3GPT execution act in concordance with each other, and the model has succeeded in combining multiple experimental representations of a pathology into a single entity that can be simulated with enough precision to carry out target identification tasks.

## Discussion

In this article, we demonstrate a novel approach to training domain-specialized language models. We have illustrated its benefits in a range of research problems, ranging from age prediction to chemical screening, with a demonstration of its utility in a real-life research project.

P3GPT shares its architecture with the open-sourced <u>MosaicML</u> platform, MPT, which is similar to LLaMA but is optimized for training and inference with linear biases instead of positional embeddings ^28,29^. Another technical solution that has led to P3GPT’s superior performance relates to omics data formatting and preprocessing.

In Geneformer, authors show that omics samples can be represented as sorted gene lists with no accompanying numerical values ^30^. We have elaborated on this concept by providing up- and down-regulated gene lists to enable a platform-agnostic and structured training process. By reducing the omics observations down to the lists of genes, data generated using practically any platform can be integrated into the framework ^30^. To ensure proper entity segregation, the omics observations reduced in this way were accompanied by metavariables (subject species, sex, tissue, etc.) flanked with tags (**Figure 5A**). This prompt format has proven to be quite useful in cross-source harmonization, since tagged fields with explicitly empty values can be imputed during inference to restore the unseen conditions of an experiment. The final aspect of the prompt structure is the instructions added to inform P3GPT of the investigators’ intent. Each instruction represents an experimental setting, such as chemical screening (<compound2diff>), case-control cross-sectional studies (<disease2diff>), and pace-of-aging studies (<age_group2diff>). This information aids P3GPT in deriving the causality inherent in the research process and offers a convenient way to control model behavior.

**Figure 5.**
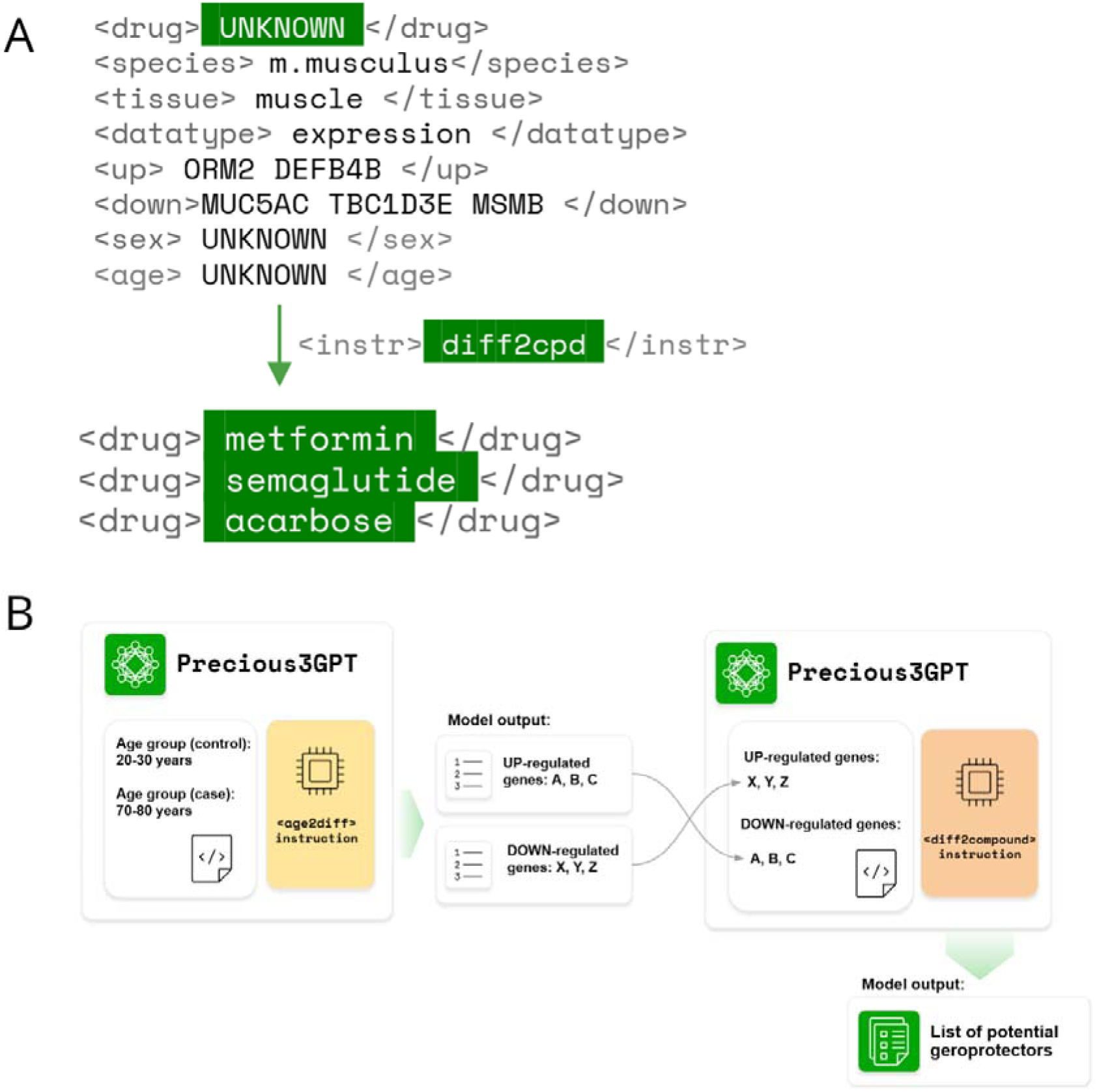
P3GPT uses a rigid prompt structure to formalize research settings and communicate with the user. **(A)** The prompt structure used by P3GPT. All possible dimensions of the tabular omics data are included in the prompt, including subject metadata. In the case of missing features, explicitly empty values are added flanked by corresponding tags (here marked as “UNKNOWN”). The prompt is supplemented with an instruction to inform P3GPT on which empty values to return based on the experimental setting. In this example, an operator uses the <diff2cpd> instruction to receive compounds that can result in the over- and under-expression of the genes included in the prompt with the <up> and <down> tags, correspondingly; **(B)** P3GPT can be used to generate novel geroprotectors. To achieve this, the model first needs to be instructed to generate the omics signatures differentiating older and younger individuals by applying the <age2diff> instruction to a prompt with the specified context. Then, the produced gene lists need to be supplied as input to P3GPT instructed to generate the compounds mimicking the reverse aging signature. Note that only age groups and gene lists are modified in the prompts throughout the workflow. Other prompt fields, such as the species, tissues, or omics type may be set to reflect a specific experimental setting.

The appropriately selected model architecture in tandem with our specialized omics data representation results in a more efficient training process requiring fewer steps to achieve domain-specific performance on par with much larger models.

Other biomedical LLMs trained on corpora of natural texts (BioBert, BioGPT, and OpenBioLLM) do not require such structured input yet still show great performance in text mining, named entity recognition, and relation extraction tasks ^4,31,32^. However, these models have a range of constraints that interfere with their adoption in research settings. Their inability to learn from multimodal data significantly limits the scope of the sources from which they can learn.

P3GPT avoids this pitfall thanks to multistep training that combines gene expression, DNA methylation, and proteomic studies with textual and KG data. This synthesis allows a broader view of biological processes, ultimately leading to better insights. Altogether, P3GPT’s design not only improves the model’s ability to understand and predict complex biological phenomena but also sets a new standard for integrating multi-omics data into LLMs.

We have demonstrated the benefits of the selected methodology in a wide range of biomedical tasks, such as age prediction, DEG prediction, target identification, and—most importantly— assessing the effects of compounds on cell lines. Surprisingly, P3GPT exhibits state-of-the-art performance in some of these tasks when compared to much larger models. Featuring only 89 mln parameters, it is measurably superior to BioGPT-Large and OpenBioLLM-8B in tests involving biological function and cell localization gene annotations (**Table3**). We view these results as an indication of the successful contextualization of biological entities within their experimental contexts. Conversely, P3GPT-derived embeddings contain less information on protein molecular function than other models do. Nonetheless, all P3GPT-based molecular function classifiers boast a ROC-AUC > 0.80, signaling the model’s adequate representation of the biochemical aspect of protein function.

We also argue that the chosen GO benchmark is a useful tool for assessing any LLM’s area of unique expertise and highlighting the range of problems to which it is applicable. In this work we only demonstrate P3GPT’s performance on a small subset of all available GO terms, although for future iterations, this benchmark should be extended to all available GO terms to provide a broader overview of a model’s capabilities.

Similarly, we have demonstrated P3GPT’s applicability in aging research by constructing a set of DNAm aging clocks that may be used in multi-species and multi-tissue settings. In the pan-mammalian clock experiment, we used the same dataset as in ^33^, which reports a clock with a Pearson’s r value of 0.96 in leave-one-species-out validation. The P3GPT-derived DNAm clock achieved a Pearson’s r value of 0.83 in the same dataset with a less robust single-species-single-clock training procedure. A direct comparison of these clocks is thus not feasible since the original pan-mammalian clock used ß-values on specific CpG sites as features that do not align with P3GPT’s prompt structure. However, this experiment allowed us to show that P3GPT’s embeddings contain aging-related information that can be applied to homologous entities in unseen species, thus implying the universality of aging mechanisms.

Moreover, we showed that P3GPT’s embeddings. This suggests that P3GPT’s multimodal approach to integrating omics data, text, and KGs offers certain benefits in tasks that require generalization across species, tissues, and omics domains.

We have also demonstrated the model’s ability to build a multi-domain view of biological phenomena as a target identification task (**Figure 4C**). Assuming that P3GPT has internalized the biological processes associated with a disease, we would expect to observe an abundance of clinically verified targets in P3GPT’s output for the <disease2diff> instruction. However, P3GPT’s output varies even if the user-defined conditions are fixed, which necessitates the generation of thousands of gene lists to meaningfully sample all the genes that may be differentially regulated in a disease. Thus, we chose to use the tokens’ output probabilities as a measure of the model-assigned importance of a gene to reduce the number of P3GPT calls to one per prompt. In total, we generated 72 sets of gene importance scores, divided across 24 diseases and 3 omics domains.

For 15 out of 24 conditions, the model successfully included clinically relevant targets among the 300 most important genes for at least one omics domain (**Figure 4C**). In 7 out of 24 cases, the most important genes were significantly enriched in clinical targets in every domain. We hypothesize that P3GPT may be well suited to driving novel target discovery under these conditions, or, more specifically, for melanoma, follicular lymphoma, chronic obstructive pulmonary disease, chronic kidney disease, and breast, pancreatic, and liver cancers (**Table S1**). Interestingly, P3GPT picks up different targets depending on the omics dimension specified in the prompt. We see this as an indication that P3GPT has successfully convoluted cross-modal representations of a pathology into a single process, thereby gaining the ability to discriminate between the different levels of regulation: epigenetic, transcriptional, and proteomic. We also hypothesize that the extracted lists of the most important genes may contain yet unknown targets and should be further investigated to assess the clinical potential of these potential targets.

For some indications, however, P3GPT fails to identify target genes as important at any level of regulation. We attribute such cases to either (i) insufficient representation of a condition in the training data or (ii) a pathogenic mechanism not represented in our model. The former may be resolved by expanding the data collection and thoughtful data preprocessing. The current iteration of P3GPT lacks the ability to assess the similarity of the two conditions. For example, EFO:0000365 (colorectal adenocarcinoma) and EFO:1001949 (colon adenocarcinoma) are two closely related indications, yet important P3GPT genes contained clinical targets only for the former (**Table S1**). The current training procedure does not explicitly convey the ontological distances between diseases, compounds, tissues, or cell lines, which likely impedes the propagation of information between similar entities.

Another major limitation imposed on P3GPT by its design is its inability to efficiently handle numeric operations. Even the most advanced LLMs are notorious for their poor performance in tasks that are not necessarily complex but require precision ^34^. Many authors have proposed ways to improve the performance of LLMs in mathematical tasks, e.g., by implementing a step-by-step solution procedure or entirely uncoupling numerical values from text token generation ^35,36^. We are planning to enable such approaches in future Precious iterations to enable quantitative analysis using synthesized data.

Despite these drawbacks, P3GPT’s performance in target identification and other tasks highlights its potential as a powerful tool for aging-related research and drug discovery. As a low-level LLM, it has the potential to enable the fast-paced, affordable experimentation setting of an *in silico* lab. In tandem with its previous and upcoming iterations, P3GPT can support a variety of complex experimental settings. To confirm this claim, we used P3GPT as a source of potential geroprotectors to be screened in a cellular senescence model (Table 4).

P3GPT allows several strategies to identify such compounds, among which we selected the method that relies on generating differential gene lists for younger and older adults (Figure 5B). Previous works that have used machine learning to identify novel senolytics have reported a 14% (3 out of 21) hit rate ^37^. In comparison, 23% of the compounds proposed by P3GPT showed senomorphic activity and no cytotoxicity. Due to time and budget constraints, we could allocate only 54 dilutions for a total of 22 compounds, resulting in < 3 dilutions per compound. We think that the actual P3GPT hit rate in geroprotector discovery could be even higher if a wider range of concentrations was explored.

For example, XL-888 was recently identified as a senolytic in an *in vitro* model of lung fibrosis ^38^. This senolytic activity is likeliest realized via HSP90 inhibition, a well-documented pathway activated by some other senolytics ^39^. We independently identified XL-888 as a potential senolytic, although all four of its dilutions displayed a significant cytotoxic effect in nonsenescent IMR90 cultures (Figure 4B). However, we also observed that, in lower concentrations, this compound is selectively more toxic to senescent cells, and we expect that, at concentrations < 20nM, it will have a negligible effect on the non-senescent cells in our *in vitro* model.

Other promising compounds selected by P3GPT for this screening include maslinic acid, a natural terpenoid found in olives, which has previously been reported to alleviate aging-related disorders in muscle and cartilage ^40^. Here, we report it to be a senomorphic compound that prevents senescent cell formation, a mechanism of action through which beneficial effects can be achieved in the elderly.

For other identified senomorphics, there is scarce evidence of geroprotective effects in the literature. Clomethiazole, for instance, is a sedative reported to have neuroprotective effects in ischemia^41^.

Dapsone was previously reported to exert an anti-inflammatory effect by inhibiting reactive oxygen species production ^42^. One compound that demonstrated potential senolytic action in our *in vitro* screening, metronidazole, has been used since the 1950s to treat inflammatory gastrointestinal conditions, such as colitis, thanks to its cytotoxic effect on anaerobic bacteria ^43^. Despite a long history of research, the metabolism of metronidazole by human cells is still not fully understood, and our findings suggest that this compound may be worthwhile to study outside the antibiotic context^44^.

While none of the screened compounds was as potent as rapamycin or navitoclax, we see this as a technical matter of identifying the optimal dosage. Alternatively, compounds with well-studied mechanisms of action, such as XL-888, may be supplemented with other potential geroprotectors to achieve a synergistic elimination of senescent cells. While P3GPT is designed to simulate experiments with singular compounds, we aim to enable the screening of compound mixtures in future Precious iterations.

In sum, P3GPT is a unique biomedical model, and its development necessitates the creation of a whole range of ancillary solutions to leverage the recent advancements in AI technology. The rapid rates of LLM releases, biodata repository updates, and research publications pose a considerable challenge to benchmarking biomedical LLM performance. Locating and aggregating the latest versions of all materials required for validation tests, as well as updating the existing pipelines to support disparate data formats, requires substantial effort that is better spent elsewhere. To reduce the burden of these routine tasks on P3GPT developers, we sought ways to automate them using AI agents.

AI agents have proven to be instrumental in overcoming common limitations of LLM applications by providing them with a toolset to actively seek new information, carry out complex pipelines, and reduce the uncertainty inherent to their output. In addition to significantly extending the range of LLM use cases, AI agents have the benefit of being mostly agnostic to the models to which they are attached, thus enabling painless transfer to newer and more powerful LLMs. Multi-agent systems have the potential to greatly increase the throughput of natural and computer science researchers without sacrificing precision, as recently demonstrated in ChatMOF, an AI system toolkit that manages demanding tasks in materials science ^45^.

The wider infrastructure that supported P3GPT’s development also featured autonomous AI agents to streamline the validation process. We have successfully enabled most tests featured in this preprint with the functionality provided by <u>Langchain</u> and <u>CrewAI</u>. As the Precious project and the broader AI field continue to mature, we intend to extend the range of functions carried out by AI agents. Ultimately, we envision a not-so-distant future in which AI systems manage end-to-end research projects in which a Precious-like model occupies a central role (**Figure 2**).

## Acknowledgements

Not applicable

## Contributions

KA — Visualization, Writing (review and editing)

AA — Supervision

FG — Writing (original draft, review and editing), Validation, Formal analysis, Visualization

VG — Writing (review and editing)

RG — Data curation (single-cell data)

AK — Data curation

SM — Data curation (single-cell data, proteome)

VN — Project administration, Supervision, Methodology (P3GPTarchitecture)

AO — Data curation (macaque data, single-cell data)

SP — Software, Investigation, Methodology (agent-based validation)

FR — Writing (review and editing)

AS — Investigation (geroprotector review)

DS — Investigation (agent-based validation), Software (mammalian aging clock), Methodology (model architecture)

QT — Investigation, Methodology (lab validation)

AT — Writing (review and editing)

AU — Investigation (agent-based validation), Data curation, Formal analysis

DX — Investigation (lab validation)

KY — Data curation (aging clocks)

DZ — Software, Data curation

AZ — Conceptualization, Supervision, Resources

## Data & code availability

P3GPT is publicly available at Insilico Medicine’s Hugging Face repository (https://huggingface.co/insilicomedicine): <u>precious3-gpt</u> (multi-omics model trained only on omics data) and <u>precious3-gpt-multi-modal</u> (multi-omics and multi-modal model) ^46,47^. Limited access to a private Hugging Face endpoint is available at Insilico Medicine’s <u>official Discord server</u> (https://discord.gg/DMAKp5yM).

Supplementary materials for this preprint are available as an Open Science Framework repository (https://doi.org/10.17605/OSF.IO/QRT3U) ^48^.

## Methods

### Data collection and processing

We aggregated omics data from experiments deposited in public repositories (see Table 2 for a full list of datasets included in the model’s training). To conform with the P3GPT-enforced prompt structure, omics measurements were reduced to lists of differentially expressed genes for RNAseq and expression profiling through array experiments, differentially methylated TSS1500 regions for RRBS and methylation profiling through array experiments, or differentially abundant proteins for proteome experiments.

For LINCS, to take one example, we used <u>level-5 data</u> representing DEG signatures as defined by the consortium. Only LINCS chemical perturbation experiments were collected for the model’s training. We defined resources that did not report results as lists of features differentiating case and control cohorts using DESeq2 for expression data ^49^. For methylation data, we used custom scripts to calculate the average []-value among all CpG sites observed in an experiment for TSS1500 regions, regardless of the platform.

Data from three species were included in P3GPT’s training: *Homo sapiens*, *Mus musculus*, and *Macaca mulatta*. For *H. sapiens*, we mapped any gene representations encountered onto NCBI gene symbols, ignoring transcript variants. Only protein-coding genes were considered, yielding a background set of 25,332 unique genes. For *M. musculus* and *M. mulatta*, all genes were denoted as the gene symbol of their human homolog with the highest similarity, as described on <u>ENSEMBL</u>. Genes that lacked human homologs were ignored.

Human clinical blood tests used in the model’s training included 30 blood biomarkers representing red blood cell indices, white blood cell indices, lipid profiles, liver proteins (ALT and AST), total bilirubin, serum minerals (calcium, phosphate, potassium, and sodium), and serum protein, albumin, and globulin levels. All blood biomarkers were included in the training procedure alongside their numeric values after min–max scaling of the original SI units.

The collected data were eventually formatted as sentences with a strictly defined structure. We represented an omics experiment as a row in a table with a predefined set of columns. Distinct columns were used for lists of upregulated and downregulated genes alongside species, data type, tissue, cell line, gender, chronological age, age group, condition, administered compound, compound dose, and duration of exposure.

Each row in the resulting table was treated as a unique sentence. Missing features were noted as explicitly empty in the sentence structure in cases of insufficient annotation in the original source or an incompatible experimental design (e.g., chemical perturbation in healthy subjects). An instruction was added at the beginning of each sentence to represent the experimental design: chemical omics screening (<cpd2diff> and <diff2cpd>), cross-age group observational study (<age2diff> and <diff2age>), or case-control condition study (<diff2disease> and <disease2diff>). In some cases, the instructions were chained to represent composite designs, e.g., <cpd2diff><disease2diff> for experiments in which a compound was applied to cancer cells. Each token, except tags, contained one white space at the end to ensure that all values were tokenized correctly.

The GEO datasets used to train the model included the following: GSE174065, GSE094274, GSE069756, GSE174196, GSE163443, GSE111234, GSE146178, GSE051264, GSE096732, GSE131194, GSE162817, GSE041637, GSE118438, GSE049379, GSE089148, GSE124709, GSE186969, GSE179330, GSE181487, GSE056845, GSE123936, GSE157690, GSE142760, GSE132496, GSE190659, GSE155443, GSE159347, GSE193264, GSE078165, GSE017274, GSE112536, GSE151815, GSE108676, GSE144783, GSE050781, GSE030352, GSE033588, GSE081382, GSE142585, GSE159214, GSE040499, GSE053260, GSE064797, GSE060269, GSE158934, GSE148290, GSE120271, GSE061420, GSE148132, GSE122044, GSE038572, GSE163177, GSE070299, GSE156161, GSE029629, GSE030198, GSE179722, GSE086939, GSE112537, GSE095736, GSE153082, GSE149758, GSE134707, GSE043520, GSE112535, and GSE072879.

### Model architecture and training

The two components of the architecture we used are the primary language model and the auxiliary modality extension (**Figure 1**).

The primary component is based on the MPT architecture, a decoder-only LLaMA-like transformer model, which was optimized for efficiency by replacing positional embeddings with a matrix of linear biases within the attention layer ^29^. The model features 32 transformer blocks with 32 self-attention heads each, as well as 256-dimension-long token embeddings, amounting to 41 million parameters. The number of parameters for P3GPT was selected based on the Chinchilla scaling law, providing a ratio of roughly 1:10 to training set tokens ^50^.

The auxiliary component enriches P3GPT with multimodal data through three-layer feed-forward neural networks (modality mappers, MM). MMs take in embeddings from additional modalities— biomedical texts and KGs—and map them onto the language model’s latent space, effectively fusing diverse knowledge sources. For the knowledge graph modality, we used embeddings generated by a heterogeneous graph transformer trained on data assembled using the Indra tool ^51–53^. For biomedical text, we utilized embeddings from Open AI’s text-embedding-ada-002 accessed via <u>GenePT</u> ^54^. Each gene’s embedding was averaged across the up- and down-regulated gene lists to create representations of gene lists. The training involved optimizing the language modeling objective using the AdamW optimizer with a dynamic learning rate, starting at 5e-3 and decaying by 0.01 after the 6th and 10th epochs, over a total of 15 epochs ^55^. The training was conducted on five A6000 GPUs with a batch size of 17.

### Agent-based validation

In our study, we employed an autonomous agent-based validation pipeline to assess P3GPT’s capabilities in various aging-related and biomedical tasks. This pipeline involves multiple stages, each designed to procure the necessary data, attach LLMs deposited on Hugging Face, extract entity embeddings, and evaluate different aspects of the model’s performance. The pipeline was realized using <u>Crew AI</u> and <u>Langchain</u> tools.

#### Gene list generation

The 805 randomly selected chemical perturbations were excluded from the LINCS training subset. The prompts for the P3GPT-derived lists used the following structure:

~~~
<disease2diff2disease><compound2diff2compound><tissue>blood</tissue> <cell>thp1</cell><efo>efo_0000221</efo> <drug>fluconazole</drug> <dose>0.125um</dose><time>24h</time> <species>human</species>
~~~

The prompts for the GPT4o-derived lists used the following structure:

Identify 250 down-regulated genes in breast mcf7 cells isolated from a human with breast adenocarcinoma following treatment with brd-k84421793. Return only genes in list separated by comma without numeration, e.g GeneA, GeneB, …. Don’t add any comments and apologies. Even if you don’t know the answer, suggest the most likely list of genes.

The generated up- and down-regulated lists were truncated to the length of the corresponding level-5 signatures. The random model was implemented as a random no-replacement sampler over the union of roughly 11,000 gene symbols present in the original LINCS holdout set.

The performance of the LLMs was measured as the Jaccard similarity of the generated DEGs to the original lists of up- and down-regulated genes. The significance of the observed differences in similarities between P3GPT, GPT4o, random sampler-derived gene lists and LINCS signatures was measured with Mann-Whitney’s U-test.

#### P3GPT entities are associated with biological processes

We manually selected six high-order terms from the three Gene Ontology (accessed on 2024-01-17) sections: molecular function, biological process, cell component. The embeddings from P3GPT (N_dimensions_ = 365), OpenBioLLM-8B (N_dim_ = 4096), BioGPT-Large (N_dim_ = 1600), Gemma-2B (N_dim_ = 2048), Llama3-8B (N_dim_ = 4096) were extracted using local instances of these LLMs installed from their Hugging Face repositories. For each gene embedding, binary prediction targets were defined as the presence of an ontology term in its annotation.

The embeddings of all genes were submitted to a CatBoost regressor training pipeline with 2000 iterations to predict whether a gene had a particular term with the following settings. The standard deviation of the reported ROC-AUC was measured based on CatBoost performance in five-fold cross-validation.

#### P3GPT-derived aging clocks

Illumina array DNAm data required for the human aging clocks was prepared following the methodology of ^56^ to obtain the total set of 10,750 samples, from which 2,824 samples were split into a validation set. The DNAm features for the reported CatBoost regressor of chronological age were selected using SHAP value feature importance scores. Among the 25,332 genes, we selected the 360 most important ones to match the length of P3GPT’s embeddings. Then, their ß-values were concatenated with the averaged embeddings of the 50 most hypermethylated genes in each sample.

The predictions for other human aging clocks were obtained from a repository of aging clocks, ClockBase ^14^. The aging clocks from the following publications were used in our comparison: ^15–19,57,58^. 618 samples present in our validation set were missing in ClockBase, hence all cross-clock comparisons were carried out in a common subset of 2,206 samples.

A CatBoost regressor was trained with the stacked embeddings of the 50 most methylated gene TSS1500 regions.

For the multi-species aging clocks, we used processed ß-values from GSE223748 GEO dataset. The samples of each species were split with a ratio of 8:1 training to test points. The cohorts with <30 points in the test set were removed. For each cohort, a separate CatBoost regressor was trained using the stacked embeddings of the 1000 most methylated gene promoters. For mapping non-human CpG sites to human gene symbols, the manifest file for GPL28271 from the original paper was used ^59^. The orthologous CpG probes were considered if they occupied TSS200 regions. The signal across several CpG sites mapped to the TSS200 region of a single gene was averaged.

#### Multimodal target enrichment

Indication-target linkage was collected by cross-referencing the mechanism of action and recorded indications for all <u>CHEMBL33</u> compounds (release date: May 2023). The level of clinical approval for a compound in the context of an indication was ignored. We selected the indications to be screened based on the high (>10) number of known clinical targets and limited our pool to the indications affecting the tissues with >1000 observations in the training set. The indications selected for screening are listed in **Supplementary Table S1**. The smallest number of targets was recorded for indication EFO:1001949 (colon adenocarcinoma) — 19 targets. The largest number of targets was recorded for MONDO:0007254 (breast cancer) — 194 targets.

For each indication, P3GPT was launched with prompts of the following format, once for each omics domain:

~~~
’[BOS]<disease2diff2disease><tissue>lung</tissue><efo>EFO_1001818</efo><dataset_type>expression</dataset_type> <gender>m</gender> <species>human</species>’
~~~

The <tissue> tag was set based on the primary tissue associated with the condition, the <gender> tag was left blank, unless a condition was strongly associated with only one gender (e.g. prostate of ovarian carcinoma).

For each launch, output token probability was collected to obtain lists of 300 most up- or down-regulated genes, for a total of 600 genes per an indication-omics generation. We inspected each such gene set for enrichment in CHEMBL-derived clinical targets using Fisher’s exact test over a contingency table with the total sum of cells equal to 22,509. See Supplementary Table ZXZ for the detected P-values and lists of genes included in P3GPT lists of most likely output tokens in each launch.

#### P3GPT-derived geroprotectors

##### P3GPT-mediated geroprotector generation

The geroprotector generation experiment was conducted using P3GPT models in a two-phase process:

1. Gene Expression Profiling:
Initially, Precious3GPT models were employed to generate lists of up-regulated and down-regulated genes for human lung tissue at the expression level. This was accomplished using the <age2diff> instruction protocol and specifying the case-control generation conditions as older adults (70-80 years) and younger adults (20-30 years), respectively;
2. Compound Identification:
Subsequently, the gene lists produced in Phase-1 were utilized as input for the P3GPT models’ <diff2compound> instruction. To induce a reversal of the age-related gene expression signature, the up-regulated and down-regulated gene lists were interchanged prior to input. The IMR90 cell line, which served as the experimental model for in vitro studies, was specified as an additional parameter in the model input for this phase.

This two-step approach facilitated the identification of potential geroprotective compounds that could theoretically counteract age-associated gene expression changes in human lung tissue. On manual review, addictive, highly toxic, and commercially unavailable compounds were removed from consideration, resulting in a list of 22 compounds to be screened in an in vitro IMR90 model of induced senescence.

##### Cell culture

Normal human lung fibroblast cell line IMR-90 was obtained from ATCC and cultured in MEM medium (Procell, cat# PM150411) supplemented with 10% fetal bovine serum (Gibco, cat# A5669701), 1% Non-Essential Amino Acids (Gibco, cat# 11140050), 1mM Sodium Pyruvate (Gibco, cat# 11360070) and 1% penicillin-streptomycin (P/S, Gibco, cat# 15140122). The cell line was routinely tested for mycoplasma contamination (Lonza, cat# LT07-710) and authenticated with short tandem repeat (STR) assays.

## Materials

All tested compounds were purchased from Medchemexpress (MCE) and dissolved in dimethyl sulfoxide (DMSO, Solarbio, cat# D8371). MCE catalog numbers for all compounds used in the screening are available in **Supplementary Table S2**.

### si**β**PIX senescence model induction

IMR-90 cell suspensions were seeded in 96-well plates at a final cell number of 2000 per well and transfected two siRNAs for βPIX gene (sißPIX; sense strand: GAAGUUAAGUUCAGCAAACAU, antisense strand: AUGUUUGCUGAACUUAACUUC) and non-targeting control (siNC; sense strand: CGUACGCGGAAUACUUCGAUU, antisense strand: AAUCGAAGUAUUCCGCGUACG) with RNAi duplex-Lipofectamine™ RNAiMAX Kit (ThermoFisher, cat 13778150) for 24 hours at 37°C, 5 % CO_2_. After 24 hours of incubation, the medium containing siRNA was replaced with a fresh medium containing diluted compounds with concentrations indicated in the figures. The treatment duration was 72 hours. Three replicates per treatment condition were performed. All siRNAs were purchased from GENScript and used at a final concentration of 10 nM.

### Senescence-associated beta-galactosidase (SA-β-gal) staining

After compound treatment, SA-β-gal staining was performed using the SA-β-gal staining kit according to the manufacturer’s protocol (Beyotime, cat# C0602). Briefly, cells were fixed using 100μL fixation buffer and incubated for 15 mins at room temperature (RT), then washed twice with 100μL PBS for 3 mins at RT and final aspiration. Subsequently, 100μL of staining solution was added to each well; plates were incubated overnight in a CO_2_-free incubator at 37°C. The next day, the staining solution was aspirated, and cells were washed with 100μL PBS. Then, 100μL of 4μg/mL Hoechst solution was added to each well, incubated for 15 mins at RT, and washed twice with PBS. The plates were sealed and scanned using ArrayScan High-Content Screening System (Thermo Fisher Scientific, CX7 LZR) in 5 channels (DAPI: Ex405/Em446/37; SA-β-gal: 590(Amber)-Brightfield, 617(Red)-Brightfield, 447(Blue)-Brightfield, 530(Green)-Brightfield) with 25 fields per well and 10×magnification (binning 2×2, 1104×1104). The size of each field is 885.54 × 885.54 μm.

### Statistical analysis

Senolytic, senomorphic, and cytotoxic potentials were formally defined using the total and ßGal positive cell counts recorded during chemical screening.

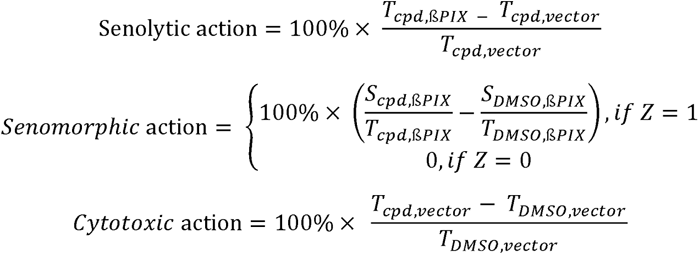

where

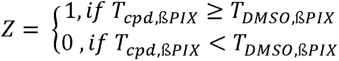

and where T_i,j_ is the total cell count in the plate wells exposed to compound *i* and transfected with siRNA *j*, and similarly, S_i,j_ is the ßGal-positive cell count.

All the aforementioned values were calculated in replicates of three for each dilution of the screened compounds. The statistical significance of non-zero action coefficients was assessed with Mann-Whithey’s U-test.

Effectively, the senolytic action of a compound represents the drop in cell survival when senescence is induced. The senomorphic action represents the decrease in the relative number of senescent cells in response to compound exposure. The senomorphic test also requires that a compound does not decrease the total cell count in the ßPIX siRNA-treated cultures, which would imply cytotoxicity and/or senolytic action. The cytotoxic action of a compound represents the decrease in cell survival when senescence is not induced.

**Table 5** reports only the dilutions with the highest achieved senolytic or senomorphic action in a compound. See **Supplementary Tables S3-6** for all compounds and dilutions tested

## Manuscript drafting

The initial draft of this text was generated using Science42: DORA (Draft Outline Research Assistant), an AI multiagent system developed by Insilico Medicine for document generation. Each agent uses Retrieval-Augmented Generation (RAG) for data collection, analysis, and fact-checking.

This LLM-based assistant is designed to streamline the process of creating publications, making it faster and simpler. The document generation process is managed by autonomous AI agents integrated with curated databases.

## Notes

https://doi.org/10.57967/hf/2699

https://doi.org/10.17605/OSF.IO/QRT3U

https://discord.gg/pSjWbmjX

### Competing Interest Statement

The authors are affiliated with Insilico Medicine, a commercial company developing and using generative artificial intelligence and other next-generation AI technologies and robotics for drug discovery, drug development, and aging research. Utilizing its generative AI platform and a range of deep aging clocks, Insilico Medicine has developed a portfolio of multiple therapeutic programs targeting fibrotic diseases, cancer, immunological diseases, and a range of age-related diseases.

## Bibliography

1. Edgar, R., Domrachev, M. & Lash, A. E. Gene Expression Omnibus: NCBI gene expression and hybridization array data repository. Nucleic Acids Res. 30, 207–210 (2002).

2. Perng, W. & Aslibekyan, S. Find the Needle in the Haystack, Then Find It Again: Replication and Validation in the ‘Omics Era. Metabolites 10, 286 (2020).

3. Vaswani, A., et al. Attention Is All You Need. Preprint at 10.48550/arXiv.1706.03762 (2023).

4. Luo, R. et al. BioGPT: Generative Pre-trained Transformer for Biomedical Text Generation and Mining. Brief. Bioinform. 23, bbac409 (2022).

5. Advancing Open-source Large Language Models in the Medical & Healthcare Domain. https://huggingface.co/blog/aaditya/openbiollm.

6. Miyakawa, T. No raw data, no science: another possible source of the reproducibility crisis. Mol. Brain 13, 24 (2020).

7. Lachmann, A. et al. Massive mining of publicly available RNA-seq data from human and mouse. Nat. Commun.9, 1366 (2018).

8. Evangelista, J. E. et al. SigCom LINCS: data and metadata search engine for a million gene expression signatures. Nucleic Acids Res. 50, W697–W709 (2022).

9. Fang, S. et al. HERB: a high-throughput experiment- and reference-guided database of traditional Chinese medicine. Nucleic Acids Res.49, D1197–D1206 (2021).

10. Kamya, P. et al. PandaOmics: An AI-Driven Platform for Therapeutic Target and Biomarker Discovery. J. Chem. Inf. Model. 64, 3961–3969 (2024).

11. Mitchell, D. C. et al. A proteome-wide atlas of drug mechanism of action. Nat. Biotechnol.41, 845–857 (2023).

12. Mamoshina, P. et al. Population Specific Biomarkers of Human Aging: A Big Data Study Using South Korean, Canadian, and Eastern European Patient Populations. J. Gerontol. Ser. *A***73**, 1482– 1490 (2018).

13. Ying, K. et al. ClockBase: a comprehensive platform for biological age profiling in human and mouse. 2023.02.28.530532 Preprint at 10.1101/2023.02.28.530532 (2023).

14. A Unified Framework for Systematic Curation and Evaluation of Aging Biomarkers | bioRxiv. https://www.biorxiv.org/content/10.1101/2023.12.02.569722v6.

15. Hannum, G. et al. Genome-wide Methylation Profiles Reveal Quantitative Views of Human Aging Rates. Mol. Cell 49, 359–367 (2013).

16. Lu, A. T. et al. DNA methylation GrimAge strongly predicts lifespan and healthspan. Aging 11, 303–327 (2019).

17. Levine, M. E. et al. An epigenetic biomarker of aging for lifespan and healthspan. Aging 10, 573– 591 (2018).

18. Lin, Q. et al. DNA methylation levels at individual age-associated CpG sites can be indicative for life expectancy. Aging 8, 394–401 (2016).

19. Belsky, D. W. et al. DunedinPACE, a DNA methylation biomarker of the pace of aging. eLife 11, e73420 (2022).

20. Haghani, A. et al. DNA methylation networks underlying mammalian traits. Science 381, eabq5693 (2023).

21. Zhang, X. et al. Identification of Bisindolylmaleimide IX as a potential agent to treat drug-resistant BCR-ABL positive leukemia. Oncotarget 7, 69945–69960 (2016).

22. Li, Z., Zhang, J., Duan, X., Zhao, G. & Zhang, M. Celastrol: A Promising Agent Fighting against Cardiovascular Diseases. Antioxid. Basel Switz. 11, 1597 (2022).

23. Lim, H. Y. et al. Celastrol in cancer therapy: Recent developments, challenges and prospects. Cancer Lett. 521, 252–267 (2021).

24. Wang, C. et al. Celastrol as an emerging anticancer agent: Current status, challenges and therapeutic strategies. Biomed. Pharmacother. Biomedecine Pharmacother. 163, 114882 (2023).

25. Gebremedhin, D., Weinberger, B., Lourim, D. & Harder, D. R. Adenosine can mediate its actions through generation of reactive oxygen species. J. Cereb. Blood Flow Metab. 30, 1777–1790 (2010).

26. Mahashur, A. et al. Pidotimod: In-depth review of current evidence. Lung India Off. Organ Indian Chest Soc. 36, 422–433 (2019).

27. Carta, S., Silvestri, M. & Rossi, G. A. Modulation of airway epithelial cell functions by Pidotimod: NF-kB cytoplasmatic expression and its nuclear translocation are associated with an increased TLR-2 expression. Ital. J. Pediatr. 39, 29 (2013).

28. Touvron, H., et al. LLaMA: Open and Efficient Foundation Language Models. Preprint at 10.48550/arXiv.2302.13971 (2023).

29. Press, O., Smith, N. A. & Lewis, M. Train Short, Test Long: Attention with Linear Biases Enables Input Length Extrapolation. Preprint at 10.48550/arXiv.2108.12409 (2022).

30. Theodoris, C. V. et al. Transfer learning enables predictions in network biology. Nature 618, 616– 624 (2023).

31. aaditya/Llama3-OpenBioLLM-70B · Hugging Face. https://huggingface.co/aaditya/Llama3-OpenBioLLM-70B.

32. Lee, J. et al. BioBERT: a pre-trained biomedical language representation model for biomedical text mining. Bioinformatics 36, 1234–1240 (2020).

33. Lu, A. T. et al. Universal DNA methylation age across mammalian tissues. *Nat*. Aging 3, 1144– 1166 (2023).

34. Bubeck, S., et al. Sparks of Artificial General Intelligence: Early experiments with GPT-4. Preprint at 10.48550/arXiv.2303.12712 (2023).

35. Zhang-Li, D., et al. Reverse That Number! Decoding Order Matters in Arithmetic Learning. Preprint at 10.48550/arXiv.2403.05845 (2024).

36. Golkar, S., et al. xVal: A Continuous Number Encoding for Large Language Models. Preprint at 10.48550/arXiv.2310.02989 (2023).

37. Smer-Barreto, V. et al. Discovery of senolytics using machine learning. Nat. Commun. 14, 3445 (2023).

38. Lee, J. Y. et al. An in vivo screening platform identifies senolytic compounds that target p16INK4a+ fibroblasts in lung fibrosis. J. Clin. Invest. 134, e173371 (2024).

39. Fuhrmann-Stroissnigg, H. et al. Identification of HSP90 inhibitors as a novel class of senolytics. Nat. Commun.8, 422 (2017).

40. Proshkina, E. et al. Terpenoids as Potential Geroprotectors. Antioxidants 9, 529 (2020).

41. Sydserff, S. G., Cross, A. J., Murray, T. K., Jones, J. A. & Green, A. R. Clomethiazole is neuroprotective in models of global and focal cerebral ischemia when infused at doses producing clinically relevant plasma concentrations. Brain Res. 862, 59–62 (2000).

42. Khalilzadeh, M. et al. A comprehensive insight into the anti-inflammatory properties of dapsone. Naunyn. Schmiedebergs Arch. Pharmacol. 395, 1509–1523 (2022).

43. Edwards, D. I. Nitroimidazole drugs--action and resistance mechanisms. I. Mechanisms of action. J. Antimicrob. Chemother.31, 9–20 (1993).

44. Dingsdag, S. A. & Hunter, N. Metronidazole: an update on metabolism, structure-cytotoxicity and resistance mechanisms. J. Antimicrob. Chemother. 73, 265–279 (2018).

45. Kang, Y. & Kim, J. ChatMOF: an artificial intelligence system for predicting and generating metal-organic frameworks using large language models. Nat. Commun. 15, 4705 (2024).

46. Insilico Medicine. precious3-gpt-multi-modal (Revision 9e240ab). (2024) doi:10.57967/hf/2699.

47. Insilico Medicine. precious3-gpt (Revision 4ca296d). (2024) doi:10.57967/hf/2700.

48. Galkin, F. Precious-3 GPT. (2024).

49. Love, M. I., Huber, W. & Anders, S. Moderated estimation of fold change and dispersion for RNA-seq data with DESeq2. Genome Biol. 15, 550 (2014).

50. Hoffmann, J., et al. Training Compute-Optimal Large Language Models. Preprint at 10.48550/arXiv.2203.15556 (2022).

51. Bachman, J. A., Gyori, B. M. & Sorger, P. K. Automated assembly of molecular mechanisms at scale from text mining and curated databases. Mol. Syst. Biol. 19, e11325 (2023).

52. Gyori, B. M. et al. From word models to executable models of signaling networks using automated assembly. Mol. Syst. Biol. 13, 954 (2017).

53. Hu, Z., Dong, Y., Wang, K. & Sun, Y. Heterogeneous Graph Transformer. Preprint at 10.48550/arXiv.2003.01332 (2020).

54. Y, C. & J, Z. GenePT: A Simple But Effective Foundation Model for Genes and Cells Built From ChatGPT. BioRxiv Prepr. Serv. Biol. (2024) doi:10.1101/2023.10.16.562533.

55. Loshchilov, I. & Hutter, F. Decoupled Weight Decay Regularization. Preprint at 10.48550/arXiv.1711.05101 (2019).

56. Fedor Galkin, Polina Mamoshina, Kirill Kochetov, Denis Sidorenko, & Alex Zhavoronkov. DeepMAge: A Methylation Aging Clock Developed with Deep Learning. Aging Dis. 0 (2020) doi:10.14336/AD.2020.1202.

57. Horvath, S. DNA methylation age of human tissues and cell types. Genome Biol.14, R115–R115 (2013).

58. Lu, A. T. et al. DNA methylation GrimAge version 2. Aging 14, 9484–9549 (2022).

59. Arneson, A. et al. A mammalian methylation array for profiling methylation levels at conserved sequences. 2021.01.07.425637 Preprint at 10.1101/2021.01.07.425637 (2021).

